# Pymportx: Facilitating Next-Generation Transcriptomics Analysis in Python

**DOI:** 10.1101/2024.07.12.598873

**Authors:** Paula Pena González, Dafne Lozano-Paredes, José Luis Rojo-Álvarez, Luis Bote-Curiel, Víctor Javier Sánchez-Arévalo Lobo

**Affiliations:** Molecular Oncology Group, Biosanitary Research Institute, Faculty of Experimental Sciences, Francisco de Vitoria University (UFV), 28223 Madrid, Spain; Pathology Department, Hospital 12 de Octubre, Av. Córdoba, s/n, 28041 Madrid, Spain; Department of Signal Theory and Communications, Universidad Rey Juan Carlos (URJC), 28942 Fuenlabrada (Madrid), Spain

**Author notes:** Both authors contributed equally (Paula Pena González, Dafne Lozano-Paredes).

**Keywords:** Data Importation, Transcriptomics, Python, Gene Expression Analysis, Open Source, Interoperability

## Abstract

The efficient importation of quantified gene expression data is pivotal in transcriptomics. Historically, the R package Tximport addressed this need by enabling seamless data integration from various quantification tools. However, the Python community lacked a corresponding tool, restricting cross-platform bioinformatics interoperability. We introduce Pymportx, a Python adaptation of Tximport, which replicates and extends the original package’s functionalities. Pymportx maintains the integrity and accuracy of gene expression data while improving processing speed and integration within the Python ecosystem. It supports new data formats and includes tools for enhanced data exploration and analysis. Available under the MIT license, Pymportx integrates smoothly with Python’s bioinformatics tools, facilitating a unified and efficient workflow across the R and Python ecosystems. This advancement not only broadens access to Python’s extensive toolset but also fosters interdisciplinary collaboration and the development of cutting-edge bioinformatics analyses.

**Availability and Implementation:** Pymportx is released as an open-source software under the MIT license. The source code is available on GitHub at https://github.com/victorsanchezarevalo/Pymportx.

## 1. Introduction

In the realm of transcriptomics research, the accurate importation and preprocessing of expression data are paramount for reliable biological interpretation. Variations in quantification methodologies, sequencing platforms, or experimental setups introduce technical biases that, if unaddressed, can significantly distort the biological conclusions drawn from such data. Numerous strategies have been developed to mitigate these issues, encompassing both classical statistical techniques, such as normalization and batch effect correction, and more recent machine learning approaches. However, many of these methods struggle with the inherent complexities of transcriptomics data, such as low sample sizes or the need to harmonize across multiple batches or studies.

The Tximport tool in R has been pivotal in addressing part of this challenge by providing a robust framework for importing and summarizing transcript quantification data from various sources (Soneson, Love, and Robinson 2016). Its approach effectively bridges the gap between raw sequencing outputs and downstream statistical analysis, but until now, a direct counterpart in Python was missing.

Here, we introduce Pymportx, a Python package that faithfully reimplements the Tximport functionality. Pymportx not only allows for the efficient importation of transcriptomics data into Python’s rich data analysis ecosystem but also enhances the process with additional features designed to handle the diverse and complex nature of such data. By facilitating a seamless transition of Tximport’s capabilities into Python, Pymportx opens new avenues for transcriptomics research that leverages Python’s advanced computational libraries.

In direct comparison with the original R package (Soneson, Love, and Robinson 2016), Pymportx demonstrates equivalent functionality in aggregating and preparing transcriptomics data for analysis, ensuring researchers can rely on the integrity and accuracy of their processed data. Furthermore, Pymportx introduces improvements in processing speed and flexibility, allowing for more efficient workflows and the integration of Python-based data analysis tools. This makes Pymportx not just a tool for data importation but a cornerstone for the next generation of transcriptomics analysis in Python, enabling researchers to tackle the complexities of biological data with unprecedented ease and efficiency.

## 2. Implementation

Pymportx adapts the core functionality of the R package Tximport for use within the Python ecosystem, focusing primarily on the accurate and efficient importation and summarization of transcript quantification data from RNA-Seq experiments. The tool is designed to handle outputs from a variety of transcript quantification software, effectively converting them into a format that is readily usable for downstream analyses, including differential expression analysis with tools like PyDESeq2 (Muzellec et al. 2023)

### 2.1 Available Features and Code Structure

Pymportx incorporates features that align with the primary capabilities of its R counterpart. The tool is structured around a core set of functions that facilitate these processes, ensuring that users can efficiently transition from data quantification to analysis within the Python environment. This achievement includes the ability to process quantification outputs into summarized count matrices, ensuring compatibility with a range of data formats from leading quantification tools such as Salmon, Kallisto, Sailfish, and RSEM. Additionally, Pymportx supports essential transformations and scaling adjustments required for data analysis. These enhancements are integrated through a core set of functions designed to streamline the transition from data quantification to analysis within the Python environment, enhancing usability and efficiency for users.

The architecture of Pymportx leverages widely used Python libraries such as Pandas for data manipulation and NumPy for numerical computations, ensuring high performance and interoperability with the broader Python data science ecosystem. By building on the foundation provided by established Python libraries and adhering to best practices in software development, Pymportx presents a robust and user-friendly tool for the bioinformatics community. Its open-source nature, under the MIT license, invites contributions from developers and researchers alike, fostering a collaborative environment for the continuous improvement and expansion of its functionalities.

In summary, Pymportx is a vital link in the chain of RNA-Seq data analysis in Python, enabling researchers to effectively bridge the gap between data quantification and expression analysis. With its focus on accuracy, efficiency, and interoperability, Pymportx sets the stage for advanced bioinformatics research conducted entirely within the Python ecosystem.

### 2.2 Comparison with Tximport

To evaluate the performance and accuracy of Pymportx, comparative tests were conducted using tests from the Tximport GitHub repository (https://github.com/thelovelab/tximport/tree/devel/tests). These tests cover a range of input scenarios, utilizing output files from three popular RNA-seq quantification tools, namely, Salmon, RSEM, and Kallisto, in which each test is structured with its own set of input files and specific parameters. The resulting matrices from these tests are the abundance, length, and counts matrices, where the abundance matrix represents the estimated abundance of each transcript or gene in Transcripts Per Million (TPM), the length matrix contains the effective lengths of transcripts or genes, and the counts matrix provides the estimated read counts for each feature. These tests were executed on the same machine using both Tximport and Pymportx. The results demonstrate that outputs from both packages are identical, confirming the accuracy of the Pymportx implementation. To quantify this similarity, correlation coefficient, Mean Absolute Error (MAE), and Mean Squared Error (MSE) were calculated between the resulting matrices from each test, all of which support the equivalence of the outputs (**Tables 1, 2, 3 and 4**,). Furthermore, the execution time for each package was measured across all tests, with each test repeated ten times to calculate the average runtime and standard deviation. The analysis comparing computational speed between Python using Pyimportx and R using Tximport across multiple panels reveals significant performance differences for certain tasks suggesting that Pyimportx is significantly faster. In addition, to assess the statistical significance of the differences in execution times, a two-tailed Welch’s t-test was performed for each test case with ten degrees of freedom (**Figure 1A, B and C**). These results provide a comprehensive comparison of the computational efficiency of both packages.

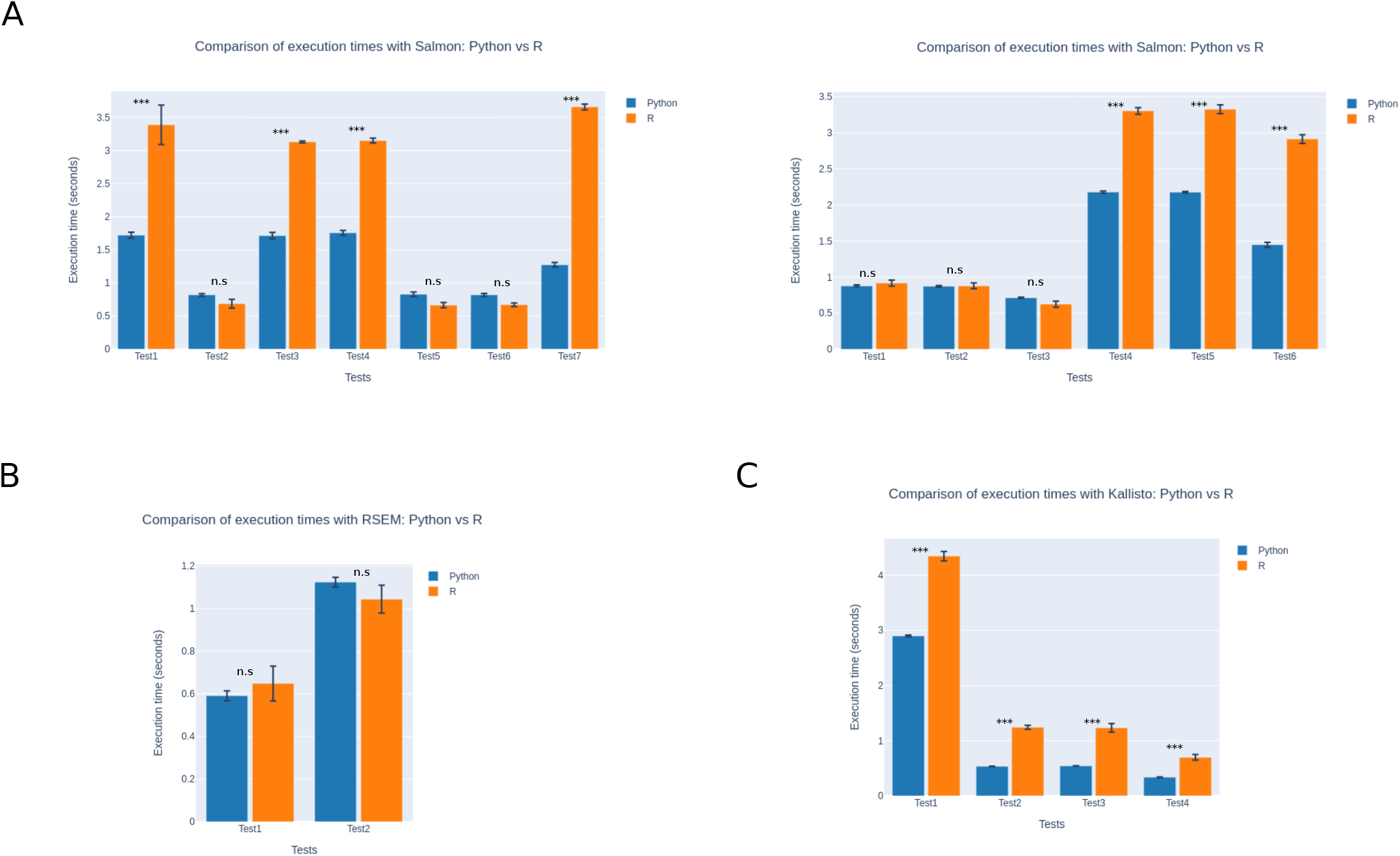

**Table 1.** Matrices resulting from the tests for the Salmon output files.

| Test | Matrix | Correlation | MAE | MSE |
| --- | --- | --- | --- | --- |
| Test 1 | Abundance | 1.0 | 5.055e-16 | 3.929e-28 |
| Test 1 | Counts | 1.0 | 1.581e-14 | 1.466e-25 |
| Test 1 | Length | 1.0 | 6.501e-13 | 2.797e-24 |
| Test 2 | Abundance | 1.0 | 0.0 | 0.0 |
| Test 2 | Counts | 1.0 | 0.0 | 0.0 |
| Test 2 | Length | 1.0 | 0.0 | 0.0 |
| Test 3 | Abundance | 1.0 | 5.055e-16 | 3.929e-28 |
| Test 3 | Counts | 1.0 | 4.175e-13 | 5.393e-23 |
| Test 3 | Length | 1.0 | 6.501e-13 | 2.797e-24 |
| Test 4 | Abundance | 1.0 | 5.055e-16 | 3.929e-28 |
| Test 4 | Counts | 1.0 | 4.522e-13 | 3.229e-23 |
| Test 4 | Length | 1.0 | 6.501e-13 | 2.797e-24 |
| Test 5 | Abundance | 1.0 | 0.0 | 0.0 |
| Test 5 | Counts | 1.0 | 1.234e-13 | 1.402e-23 |
| Test 5 | Length | 1.0 | 0.0 | 0.0 |
| Test 6 | Abundance | 1.0 | 0.0 | 0.0 |
| Test 6 | Counts | 1.0 | 1.300e-13 | 7.912e-24 |
| Test 6 | Length | 1.0 | 0.0 | 0.0 |
| Test 7 | Abundance | 1.0 | 0.0 | 0.0 |
| Test 7 | Counts | 1.0 | 1.281e-13 | 1.177e-23 |
| Test 7 | Length | 1.0 | 0.0 | 0.0 |

**Table 2.** Matrices resulting from the tests for the Salmon output files with replicates.

| Test | Matrix | Correlation | MAE | MSE |
| --- | --- | --- | --- | --- |
| Test 1 | Abundance | 1.0 | 0.0 | 0.0 |
| Test 1 | Counts | 1.0 | 0.0 | 0.0 |
| Test 1 | Length | 1.0 | 0.0 | 0.0 |
| Test 2 | Abundance | 1.0 | 0.0 | 0.0 |
| Test 2 | Counts | 1.0 | 0.0 | 0.0 |
| Test 2 | Length | 1.0 | 0.0 | 0.0 |
| Test 3 | Abundance | 1.0 | 0.0 | 0.0 |
| Test 3 | Counts | 1.0 | 0.0 | 0.0 |
| Test 3 | Length | 1.0 | 0.0 | 0.0 |
| Test 4 | Abundance | 1.0 | 8.375e-16 | 6.587e-28 |
| Test 4 | Counts | 1.0 | 2.676e-14 | 5.122e-25 |
| Test 4 | Length | 1.0 | 1.109e-12 | 4.918e-24 |
| Test 5 | Abundance | 1.0 | 8.375e-16 | 6.587e-28 |
| Test 5 | Counts | 1.0 | 2.676e-14 | 5.122e-25 |
| Test 5 | Length | 1.0 | 1.109e-12 | 4.918e-24 |
| Test 6 | Abundance | 1.0 | 8.375e-16 | 6.587e-28 |
| Test 6 | Counts | 1.0 | 2.676e-14 | 5.122e-25 |
| Test 6 | Length | 1.0 | 1.109e-12 | 4.918e-24 |

**Table 3.** Matrices resulting from the tests for the RSEM output files.

| Test | Matrix | Correlation | MAE | MSE |
| --- | --- | --- | --- | --- |
| Test 1 | Abundance | 1.0 | 0.0 | 0.0 |
| Test 1 | Counts | 1.0 | 0.0 | 0.0 |
| Test 1 | Length | 1.0 | 0.0 | 0.0 |
| Test 2 | Abundance | 1.0 | 0.0 | 0.0 |
| Test 2 | Counts | 1.0 | 0.0 | 0.0 |
| Test 2 | Length | 1.0 | 0.033 | 0.033 |

**Table 4.** Matrices resulting from the tests for the Kallisto output files.

| Test | Matrix | Correlation | MAE | MSE |
| --- | --- | --- | --- | --- |
| Test 1 | Abundance | 1.0 | 0.0 | 0.0 |
| Test 1 | Counts | 1.0 | 0.0 | 0.0 |
| Test 1 | Length | 1.0 | 0.0 | 0.0 |
| Test 2 | Abundance | 1.0 | 0.0 | 0.0 |
| Test 2 | Counts | 1.0 | 0.0 | 0.0 |
| Test 2 | Length | 1.0 | 0.0 | 0.0 |
| Test 3 | Abundance | 1.0 | 0.0 | 0.0 |
| Test 3 | Counts | 1.0 | 0.0 | 0.0 |
| Test 3 | Length | 1.0 | 0.0 | 0.0 |
| Test 4 | Abundance | 1.0 | 0.0 | 0.0 |
| Test 4 | Counts | 1.0 | 0.0 | 0.0 |
| Test 4 | Length | 1.0 | 0.0 | 0.0 |

This thorough comparison provides a robust assessment of Pymportx accuracy and performance relative to its R counterpart, offering valuable insights into its efficacy as a Python-based alternative for transcript-level data analysis. The combination of output validation and performance benchmarking demonstrates the reliability and potential advantages of Pymportx in the field of bioinformatics data processing.

## 3. Discussion and Conclusion

In this paper, we introduced Pymportx, a Python library that serves as a direct analog to the R package Tximport, designed for the streamlined importation and summarization of transcriptomics data (Soneson, Love, and Robinson 2016). This tool marks a significant step towards creating a cohesive ecosystem for transcriptomics analysis in Python. Notably, Pymportx facilitates a seamless linkage with PyDESeq2 (Muzellec et al. 2023), a Python adaptation of the R package DESeq2 (Love, Huber, and Anders 2014), which is renowned for its capabilities in differential gene expression analysis. This integration is pivotal for researchers seeking to maintain a continuous workflow entirely within Python, especially those utilizing quantification outputs from tools like Salmon and Kallisto (Patro et al. 2017; Bray et al. 2016).

The advent of Pymportx extends far beyond mere data importation; it is about enabling researchers to harness the full power of Python for transcriptomics analysis. By ensuring compatibility with PyDESeq2, we bridge the gap between data preprocessing and differential expression analysis, allowing for a streamlined, efficient workflow that leverages Python’s computational prowess and its rich ecosystem of data analysis tools. This continuity is critical for facilitating sophisticated analyses without the need to oscillate between programming languages, thus optimizing both the analytical process and the utilization of computational resources.

The open-source nature of Pymportx, under the MIT license, underscores our vision of collaborative advancement in the bioinformatics field. We encourage the community’s engagement with Pymportx, aiming for enhancements, extensions, and innovations that will drive the tool forward. This collaborative spirit is integral to the development of a robust, dynamic ecosystem of bioinformatics tools in Python.

Python’s ascendancy as a general-purpose programming language, coupled with its extensive application in data science, machine learning, and other computational fields, positions it as an ideal platform for bioinformatics. Through tools like Pymportx, we not only facilitate this transition but also enrich the analytical capabilities available to researchers, empowering them to achieve more comprehensive, nuanced biological insights.

In the era of large language models and automated code generation, the development of Pymportx serves as a beacon for the potential to expedite the porting of essential bioinformatics tools to Python, while emphasizing the indispensable role of human expertise in ensuring accuracy and optimizing performance.

As we look forward to the continued evolution of Pymportx, our commitment to open-source principles and community collaboration remains steadfast. By fostering an environment of shared knowledge and collective innovation, we anticipate that Pymportx will play a crucial role in advancing the Python bioinformatics ecosystem, paving the way for a future where complex, multi-stage analyses are conducted with unprecedented efficiency and precision.

## Acknowledgements

The authors express gratitude to Jose Luis Rodriguez Peralto, Maite Iglesias Badiola, Cruz Santos Tejedor and Alberto Lopez Rosado for their unwavering support to the group.

## Author contributions

PP, DL, LB and VJSAL implemented Pymportx. LB and VJSAL wrote this manuscript. JLR reviewed code and provided scientific guidance. All authors read and approved the final manuscript.

